# Forensic features and genetic legacy of the Baloch population of Pakistan and the Hazara population across Durand-line revealed by Y chromosomal STRs

**DOI:** 10.1101/2020.11.21.392456

**Authors:** Atif Adnan, Shao-Qing Wen, Allah Rakha, Rashed Alghafri, Shahid Nazir, Muhammad Rehman, Chuan-Chao Wang, Jie Lu

## Abstract

Hazara population across Durand-line has experienced extensive interaction with Central Asian and East Asian populations. Hazara individuals have typical Mongolian facial appearances and they called themselves descendants of Genghis Khan’s army. The people who speak the Balochi language are called Baloch. Previously, a worldwide analysis of Y-chromosomal haplotype diversity for rapidly mutating (RM) Y-STRs and with PowerPlex Y23 System (Promega Corporation Madison, USA) kit was created with collaborative efforts, but Baloch and Hazara population from Pakistan and Hazara population from Afghanistan were missing. A limited data with limited number of markers and samples is available which poorly define these populations. So, in the current study, Yfiler Plus PCR Amplification Kit loci were examined in 260 unrelated Hazara individuals from Afghanistan, 153 Hazara individuals, and 111 Balochi individuals from Baluchistan Pakistan. For the Hazara population from Afghanistan and Pakistan overall, 380 different haplotypes were observed on these 27 Y-STR loci, gene diversities ranged from 0.51288 (DYS389I) to 0.9257 (DYF387S1) and haplotype diversity was 0.9992 +/- 0.0004. For the Baloch population, every individual was unique at 27 Y-STR loci, gene diversity ranged from 0.5718 (DYS460) to 0.9371(DYF387S1). Twelve haplotypes shared between 178 individuals while only two haplotypes among these twelve were shared between 87 individuals in Hazara populations. Rst and Fst pairwise genetic distance analyses, multidimensional scaling (MDS) plot, Neighbor-joining (NJ) tree, linear discriminatory analysis (LDA), and median-joining network (MJNs) were performed, which shed light on the history of Hazara and Baloch populations. Interestingly null alleles were observed at DYS448 with specific mutation patterns in Hazara populations. The results of our study showed that the Yfiler Plus PCR Amplification Kit marker set provided substantially stronger discriminatory power in the Baloch population of Pakistan and the Hazara population across the Durand-line.

## INTRODUCTION

The variation pattern in Human DNA usually provides a balance between natural selection and neutral processes. Y chromosomal variant analysis for determining the patterns of present and past flow of genes between populations is very helpful ^1^. Y-chromosome short tandem repeats (YSTRs) plays an important role in forensic molecular biology ^2–5^. Usually, Y-STRs are used for (i) decidedly determine the male component of DNA mixtures under the presence of a high female DNA background as typically confronted with materials from sexual assault cases^6^, (ii) to test for paternal relationships between male individuals particularly in deficiency paternity cases with the mother not being available^7^, or (iii) for special cases in missing-person or (iv) disaster-victim identification involving males^8^, or (v) for evolutionary purposes because male family members share same haplotype distribution which may be different from individual to individual within a population group, or (vi) different geographic regions or in different ethnic groups. Normally, more paternal lineages can be differentiated with an increased number of Y-STRs ^9^, such as the Powerplex Y Kit (Promega) containing 12 Y-STRs ^10^, the AmpFlSTR Y-filer PCR Amplification Kit (Life Technologies) (subsequently referred to as Y-filer) containing 17 Y-STRs ^11^ or Powerplex Y23 Kit (Promega) containing 23 Y-STRs ^12^, relative to the initially proposed 9-loci haplotype ^13^. So, Applied Biosystems have developed Yfiler Plus PCR Amplification Kit ^14^. The Yfiler Plus kit provides enhanced discrimination power because it includes the Yfiler loci and 10 additional STRs in which 6 are rapidly mutating (RM) Y STRs. These rapidly mutating Y STRs showed a higher mutation rate of about a few mutations every 100 generations per locus (μ > 10^-2^) compared with all other commonly used Y-STRs. Molecular biological and cytogenetical studies give us an insight into the presence of many structural variants within the human Y chromosome, which might be deletions^15–17^, duplications^18–20^, and inversions ^19–23^. Null alleles or allele droop-out are well-established factors that can occur with any PCR-based STR typing system. The reason could be the primer binding site problem or deletions within the target region ^24,25^. DYS448 lied in the proximal part of the azoospermia factor c (AZFc) region, which is considered important in spermatogenesis and made up of “ampliconic” repeats which act as substrates for nonallelic homologous recombination (NAHR). NAHR could delete larger blocks of the Y chromosome which included DYS448^26^. This null alleles or allelic drop-out phenomenon is more commonly observed in Central Asian and East Asian populations but in the Hazara population of Pakistan, its occurrence was >16% ^27^.

Durand Line is a boundary established in the Hindu Kush around 1893 running through the tribal lands between Afghanistan and British India (modern-day Pakistan), marking their respective scopes of influence. The recognition of this line, which was named after Sir Mortimer Durand, has settled the Indo-Afghan frontier problem for the rest of the British period. Now, this is an established border between Afghanistan and Pakistan. The origin of the Hazara population is disputed. The Hazara could be of Turko-Mongol ancestry and theorized to be the descendants of an occupying army left in Afghanistan by Genghis Khan in thirteen hundred AD^28^. The Hazara population speaks Persian with some Mongolian words. The total population of Hazaras in the world is 4.5 million. Afghanistan is considered the mainland for the Hazara population (3 million) and they are the third largest ethnic group (9%) after Tajiks (27%) and Pashtuns (42%) ^29^, while in Pakistan, Hazara is one of the distinct but small groups comprising 0.08% of the total population (http://www.pbscensus.gov.pk). The tribes who speak the Balochi language are called Baloch^30^. Balochi population is 3.6% of total Pakistani population (http://www.pbscensus.gov.pk). They are also found in the neighboring areas of Iran and Afghanistan. Perhaps, the origin of Baloch homeland lay on the Iranian plateau. The Baloch were mentioned in Arabic chronicles of the 10th century. The Seljuq invasion of Kermān in the 11th century started the eastward migration of the Balochi population^30^.

In this study, we have investigated the Baloch and Hazara population from Pakistan and the Hazara population from Afghanistan using 27 Y STRs to determine their genetic history and gene diversity. This data has defined the Hazara and Baloch populations better and are supplement to the Y STR haplotype reference database (YHRD).

## 2. RESULTS AND DISCUSSIONS

### 2.1 Allelic frequencies and Forensic parameters

We successfully obtained genotypes of 524 individuals in three ethnic groups (Balochi population, Hazara population from Afghanistan, and Pakistan) (**Supplementary Table 1**). Allelic frequencies of Baloch ethnic group from Baluchistan, Pakistan, and Hazara ethnic groups from Pakistan and Afghanistan along with gene diversity values were shown in **Supplementary Table 2**.

DYF387S1 showed the highest gene diversity/heterozygosity in Baloch and both Hazara populations from Afghanistan and Pakistan with 0.9371, 0.9242, and 0.8792, respectively. Overall DYS570 (0.8624) showed the highest or DYS437 (0.2383) showed the lowest gene diversity/heterozygosity for single Y STR markers. Within three populations, single Y-STR markers DYS570 (0.8624), DYS449 (0.8468), DYS627 (0.7949) showed the highest gene diversity/heterozygosities while DYS460 (0.5718), DYS391 (0.3916), and DYS437 (0.2383) showed the lowest gene diversity/heterozygosities in the Baloch and both the Hazara populations from Afghanistan and Pakistan, respectively. After pooling Hazara populations together DYF387S1, DYS437 showed the highest or lowest gene diversity/heterozygosities with 0.9257 and 0.4053 respectively. The observed numbers of alleles were 222, 240, and 188 for Baloch and both the Hazara populations from Afghanistan and Pakistan, respectively on 27 Y STRs.

Allelic frequencies ranged from 0.0090 to 0.6036 in the Baloch population, 0.0038 to 0.6654 in the Hazara population from Afghanistan, and 0.0065 to 0.8627 in the Pakistani Hazara population.

We evaluated forensic parameters at seven levels (**Table 2**), the minimal 9 Y-STRs loci (DYS19, DYS389I, DYS389II, DYS390, DYS391, DYS392, DYS393, and DYS385a/b), the extended 11 Y-STRs loci (MHT+DYS438 and DYS439), PowerPlex Y12 STRs loci (extended 11 Y STRs + DYS437), Y-filer 17 STRs loci (PPY12+DYS448, DYS456, DYS458, DYS635, and Y_GATA_H4), Y21STRs loci(Y-filer + DYS481, DYS533, DYS570, and DYS576), Y27 Yfiler Plus loci (21 STRs + DYF387S1, DYS449, DYS460, DYS518, and DYS627), and 6 rapidly mutating Y STRs loci (DYS570, DYS576, DYF387S1, DYS449, DYS518, and DYS627) which are summarized in Table 2. The discrimination capacity (DC) ranged from 87.38% (the minimal 9 Y-STRs loci) to 100% (Y27 Yfiler Plus loci) with random matching probability from 0.0162 (MHT) to 0.009 (Y27 Yfiler Plus loci) and haplotype diversity (HD) ranged 0.9928 (the minimal 9 Y-STRs loci) to 1.0 (Y27 Yfiler Plus loci) in the Baloch population of Pakistan. The discrimination capacity (DC) ranged from 47.06% (the minimal 9 Y-STRs loci) to 99.35% (Y27 Yfiler Plus loci) with random matching probability from 0.0745 (MHT) to 0.0066 (Y27 Yfiler Plus loci) and haplotype diversity (HD) ranged from 0.9316 (the minimal 9 Y-STRs loci) to 0.9999 (Y27 Yfiler Plus loci) in Pakistani Hazara population while DC ranged 41.15% (the minimal 9 Y-STRs loci) to 88.46% (Y27 Yfiler Plus loci) with random matching probability from 0.0329 (MHT) to 0.0057 (Y27 Yfiler Plus loci) and HD ranged from 0.9708 (the minimal 9 Y-STRs loci) to 0.9937 (Y27 Yfiler Plus loci) for Hazara population from Afghanistan. Pooling both populations together DC ranged 40.19% (the minimal 9 Y-STRs loci) to 92% (Y27 Yfiler Plus loci) with random matching probability from 0.0334 (MHT) to 0.0032 (Y27 Yfiler Plus loci) and HD ranged from 0.9689 (the minimal 9 Y-STRs loci) to 0.9992 (Y27 Yfiler Plus loci). Interestingly six rapidly mutating Y STRs which are included in Yfiler plus kit detects high haplotype diversity (Table 2). We have observed 101 (90.99%) different haplotypes out of 111, among them, 95 (85.58%) were unique in the Baloch population and we have observed 139 (90.84%) different haplotypes out of 153, among them 131 (85.62%) were unique in Pakistani Hazara population while in Afghani Hazara population observed haplotypes were 188 (72.30%) out of 260, among them 152(58.46%) were unique. These six STRs (RM Y STRs) showed the almost same diversity, shown by PPY 23 loci. The above results are showing that Yfiler plus kit loci showed strong discrimination capacity, haplotype diversity, and random mating probabilities which provide utility for forensic identification and paternity testing in three ethnic groups (Baloch and Hazara from Pakistan while Hazara from Afghanistan).

**Table 1:** Reference Populations from Central, Eastern and South Asia populations selected as reference populations used in LDA, NJ tree and multidimensional scaling (MDS) analysis.

| No | Population | Haplotypes | Country |
| --- | --- | --- | --- |
| 1 | Afghan | 152 | Afghanistan<sup>64</sup> |
| 2 | Pathan | 125 | Afghanistan<sup>65</sup> |
| 3 | Pathan | 44 | North Afghanistan<sup>66</sup> |
| 4 | Pathan | 142 | South Afghanistan<sup>66</sup> |
| 5 | Farsi, Arab | 35 | Ahvaz, Iran (accession no YA004581) |
| 6 | Farsi, Azerbaijani | 34 | Hamedan, Iran (accession no YA004582) |
| 7 | Han | 934 | Beijing, China<sup>67,68</sup> |
| 8 | Kazakh | 231 | Xinjiang, China<sup>69,70</sup> |
| 9 | Mongol | 272 | Hulun Buir, China<sup>71</sup> |
| 10 | Mongol | 454 | Inner Mongolia, China<sup>72-74</sup> |
| 11 | Uighur | 732 | Xinjiang, China<sup>75</sup> |
| 12 | Arab | 33 | Ahvaz, Iran<sup>76</sup> |
| 13 | Azerbaijani | 39 | Tabriz, Iran (accession no YA004586) |
| 14 | Azerbaijani | 50 | Urmia, Iran (accession no YA004587) |
| 15 | Bakhtiari | 45 | Izeh, Iran<sup>76</sup> |
| 16 | Baloch | 19 | Balochistan, Iran (accession no YA003794) |
| 17 | Baloch | 59 | Zahedan, Iran (accession no YA004238) |
| 18 | Farsi | 286 | Tehran, Iran (accession no YA004580) |
| 19 | Gilak | 98 | Gilan, Iran<sup>77</sup> |
| 20 | Gilaki | 42 | Rasht, Iran<sup>76</sup> |
| 21 | Iranian | 27 | Birjand, Iran (accession no YA003902) |
| 22 | Iranian | 152 | Central Iran, Iran (accession no YA003782) |
| 23 | Iranian | 106 | Fars, Iran (accession no YA004229) |
| 24 | Iranian | 106 | Golestan, Iran<sup>78</sup> |
| 25 | Iranian | 94 | Iran (accession no YA004237) |
| 26 | Iranian | 161 | Isfahan, Iran<sup>79</sup> |
| 27 | Iranian | 127 | Mashhad, Iran (accession no YA003903) |
| 28 | Kurd | 51 | Iran (accession no YA004244) |
| 29 | Kurdish | 77 | Kermanshah, Iran (accession no YA004584) |
| 30 | Kurdish | 73 | Kurdistan, Iran (accession nos YA003795 and YA004585) |
| 31 | Lor | 37 | Lorestan, Iran (accession nos YA003796 and YA004243) |
| 32 | Lurs | 9 | Kohgiluyeh-Buyer Ahmad, Iran (accession no YA003797) |
| 33 | Mazandarani | 44 | Sari, Iran<sup>76</sup> |
| 34 | Mazani | 126 | Mazan daran, Iran<sup>77</sup> |
| 35 | Parsee | 17 | Fars, Iran (accession no YA003798) |
| 36 | Qashqae | 15 | Fars, Iran (accession no YA003799) |
| 37 | Sistani | 64 | Zabol, Iran (accession no YA004241) |
| 38 | Talysh | 15 | Masal, Iran<sup>76</sup> |
| 39 | Turk | 11 | Ardabil, Iran (accession no YA004240) |
| 40 | Zoroastrian | 6 | Yazd, Iran (accession no YA003800) |
| 41 | Luri | 60 | Ilam, Iran Kurdish (accession no YA004583) |
| 42 | Arain | 85 | Punjab, Pakistan<sup>80</sup> |
| 43 | Baloch | 98 | Balochistan, Pakistan (accession no YA004595) |
| 44 | Gujjar | 20 | Swat and Dir District, Pakistan<sup>81</sup> |
| 45 | Hazara | 160 | Balochistan, Pakistan<sup>27</sup> |
| 46 | Kashmiri | 175 | Azad Kashmir, Pakistan<sup>82</sup> |
| 47 | Kohistani | 20 | Swat and Dir District, Pakistan<sup>81</sup> |
| 48 | Pathan | 269 | Pakistan<sup>83</sup> |
| 49 | Punjabi | 383 | Punjab, Pakistan<sup>3</sup> |
| 50 | Roma | 278 | Punjab, Pakistan (accession no YA004554) |
| 51 | Saraiki | 51 | Southern Punjab, Pakistan (accession no YA004225) |
| 52 | Saraki | 148 | Punjab, Pakistan<sup>84</sup> |
| 53 | Sindhi | 98 | Sindh, Pakistan<sup>85</sup> |
| 54 | Yousafzai Pathan | 71 | KhyberPakhtunkhwa, Pakistan (accession no YA003748) |
| 55 | Tharklani, Pashtun | 20 | Swat and Dir District, Pakistan<sup>81</sup> |
| 56 | Uthmankheil, Pashtun | 20 | Swat and Dir District, Pakistan<sup>81</sup> |
| 57 | Yousafzai, Pashtun | 20 | Swat and Dir District, Pakistan<sup>81</sup> |
| 58 | Urdu, Punjabi | 241 | Punjab, Pakistan (accession no YA004381) |
| 59 | Kazakh | 305 | Kazakhstan<sup>86</sup> |
| 60 | Kazakh | 67 | East Kazakhstan, Kazakhstan (accession no YA003700) |
| 61 | Kazakh | 99 | South Kazakhstan, Kazakhstan (accession no YA003729) |
| 62 | Southwest Asian | 493 | United Kingdom<sup>87</sup> |
| 63 | British Pakistani | 132 | United Kingdom<sup>88</sup> |
| **Total haplotypes** |  | **8457** |  |

**Table 2:** Forensic parameters on 7 different levels in three ethnic groups

| **Hazara Pakistan** | **MHT 9 Y-STRs** | **EHT 11 Y-STRs** | **PPY-12 Y-STRs** | **Yfiler 17 Y-STRs** | **PPY23 21 Y-STRs** | **Yfiler plus 27 Y-STRs** | **6 RM Y STRs** |
| --- | --- | --- | --- | --- | --- | --- | --- |
| **No of Samples** | **153** | **153** | **153** | **153** | **153** | **153** | **153** |
| RMP | 0.0745 | 0.0577 | 0.0577 | 0.0123 | 0.0084 | 0.0066 | 0.0091 |
| HD | 0.9316 | 0.9485 | 0.9485 | 0.9942 | 0.9981 | 0.9999 | 0.9974 |
| No of haplotypes | 72 | 81 | 81 | 117 | 140 | 152 | 139 |
| NUH | 54 | 63 | 63 | 97 | 132 | 151 | 131 |
| DC | 0.4705 | 0.5294 | 0.5294 | 0.7647 | 0.9150 | 0.9934 | 0.9084 |
| % of Unique Haplotypes | 0.3529 | 0.4176 | 0.4176 | 0.6339 | 0.8627 | 0.9869 | 0.8562 |
| **Hazara Afghanistan** |  |  |  |  |  |  |  |
| **No of Samples** | **260** | **260** | **260** | **260** | **260** | **260** | **260** |
| RMP | 0.0329 | 0.0285 | 0.0272 | 0.0184 | 0.0129 | 0.0057 | 0.0101 |
| HD | 0.9708 | 0.9753 | 0.9765 | 0.9854 | 0.9909 | 0.9982 | 0.9937 |
| No of haplotypes | 107 | 122 | 124 | 166 | 190 | 230 | 188 |
| NUH | 64 | 81 | 83 | 132 | 157 | 207 | 152 |
| DC | 0.4115 | 0.4692 | 0.4769 | 0.6384 | 0.7307 | 0.8846 | 0.723 |
| % of Unique haplotypes | 0.2461 | 0.3115 | 0.3192 | 0.5076 | 0.6038 | 0.7961 | 0.5846 |
| **Pak-Afg Hazara** |  |  |  |  |  |  |  |
| **No of Samples** | **413** | **413** | **413** | **413** | **413** | **413** | **413** |
| RMP | 0.0334 | 0.0268 | 0.0262 | 0.0113 | 0.007 | 0.0032 | 0.0058 |
| HD | 0.9689 | 0.9756 | 0.9761 | 0.9911 | 0.9954 | 0.9992 | 0.9966 |
| No. of Haplotypes | 166 | 191 | 193 | 273 | 317 | 380 | 320 |
| NUH | 109 | 137 | 139 | 223 | 274 | 357 | 274 |
| DC | 0.4019 | 0.4624 | 0.4673 | 0.661 | 0.7675 | 0.92 | 0.7748 |
| % of Unique Haplotypes | 0.2639 | 0.3317 | 0.3365 | 0.5399 | 0.6634 | 0.8644 | 0.6634 |
| **Baloch Pakistan** |  |  |  |  |  |  |  |
| **No of Samples** | **111** | **111** | **111** | **111** | **111** | **111** | **111** |
| RMP | 0.0162 | 0.0136 | 0.0136 | 0.0095 | 0.0092 | 0.009 | 0.0114 |
| HD | 0.9928 | 0.9954 | 0.9954 | 0.9995 | 0.9998 | 1 | 0.9975 |
| No of haplotypes | 97 | 100 | 100 | 108 | 110 | 111 | 101 |
| NUH | 93 | 96 | 96 | 105 | 109 | 111 | 95 |
| DC | 0.8738 | 0.9009 | 0.9009 | 0.9729 | 0.9909 | 1 | 0.9099 |
| % of Unique haplotypes | 0.83783 | 0.8648 | 0.8648 | 0.9459 | 0.9819 | 1 | 0.8558 |

RMP= Random Matching Probability ; HD= Haplotype Diversity ; NUH= No. of unique haplotypes; DC = Discrimination Capacity

### 2.2 Phylogenetic analyses and Population comparisons

Since the anthropological or ethno-historical relationships between studied populations and reference populations which are included for analysis were already known, so we used two different methods on the basis of their similarity with *a priori* expectations. *Fst* is a standardized variance of haplotype frequency and assumes genetic drift as being the agent that differentiates populations. *Rst* is a standardized variance of haplotype size and takes into account both drift and mutation as causes of population differentiation, assuming a stepwise model in which each mutation creates a new allele either by adding or deleting a single repeat unit. To assess the relationship between these three populations (Baloch, Hazara from Pakistan and Afghani Hazaras), and the other relevant populations which are summarized in **Table 1,** pair-wise genetic distances (Rst and Fst) and their corresponding p-values were calculated and were shown in **Supplementary Table 3**. These Rst and Fst values were visualized using hierarchical clustering heat-map (**Supplementary Figure 1 a & b**). Dendrograms give us a clear picture about the organization of the data which can be compared with NJ trees or MDS plots. The utlization of mean-linkage dendrograms to Y STR data gives us a consistent basis of comparison. Heat-map matrix based on Rst values showed that Hazara from Pakistan were clustered more closely to Central and East Asian (i.e. Kazakh and Mongols) populations while the Baloch population was clustered with other Pakistani (i.e. Pathan and Sindhi) populations and Hazara from Afghanistan were clustered with local Afghan populations. On another hand, the heat-map matrix based on Fst values showed that the Hazara population from Pakistan was tightly clustered with local (i.e. Baloch, Arain, and Pathans) populations while the Hazara population from Afghanistan was clustered with Afghanistan Pathan and Northern Talysh population. The observed pattern of inter-population diversity from Rst was in support of anthropological knowledge, while that based on Fst revealed unexpected and unconvincing population affinities. These results are consistent with our previous study results ^31^. The pairwise Rst genetic distances values between Baloch and other relevant populations ranged from −0.0402 to 0.1417. According to Rst values, the Baloch population of Pakistan showed the closest genetic distance to Turks (−0.0402) from Ardabil, Iran while Kazakh (0.1417) from Gansu, China showed the greatest genetic distance. For the Afghan Hazara population, the Afghan population (0.0009) from Afghanistan showed the closest genetic distance and for the Pakistani Hazara group, the Afghan population (0.0381) from Afghanistan showed the closest genetic distance To investigate the paternal relationship among these three and other reference populations, we have generated the MDS plot (figure 1) based on pairwise Rst matrix from supplementary table 3. In the MDS plot, we have seen that the Hazara population from Afghanistan is located closer to the Afghan population from Afghanistan and the Pathan population from northern Afghanistan which is similar to the results of another study ^32^, while Pakistani Hazara lined closer to Kazakh and Mongolian population which is similar to our previous study’s results^27,33^.

**Figure 1:**
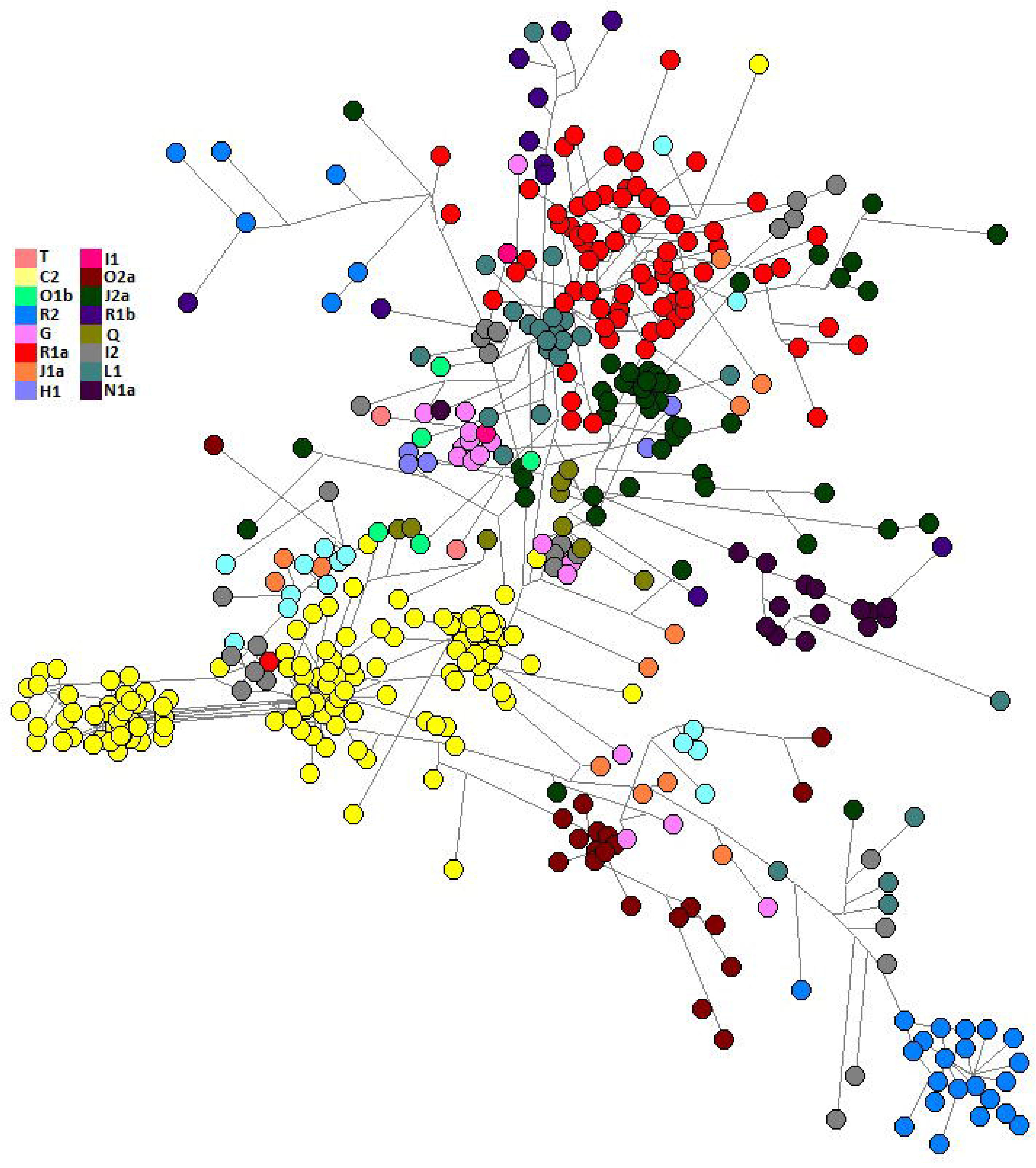
Two-dimensional plot from multi-dimensional scaling analysis of R*st*-values based on Yfiler haplotypes for the Baloch population of Pakistan and Hazara populations across the Durand line with reference populations.

According to Fst values, the Afghan Hazara population is closest to the Afghan population (0.0053) followed by the Hazara population from Balochistan, Pakistan (0.0057), and Iranian population from Mashhad, Iran (0.0077). Evolutionary relationships between the Baloch and Hazara population of Pakistan, the Hazara population from Afghanistan, and other reference populations were inferred from the Neighbor-joining tree based on Fst values (**Figure 2**). In neighbor-joining trees, an admixed population will always lie on the path between the source populations^34^. In total, we have observed 14 clusters for 62 populations in NJ-tree and the Baloch population placed itself in the second cluster along with West-south Asian populations. Hazara populations from Pakistan and Afghanistan came to the fourth cluster along with the Afghani and Iranian populations. The pattern of inter-population diversity based on *Rst* was consistent with ethnohistorical and anthropological knowledge, while that based on *Fst* shown surprising and unaccepted population affinities.

**Figure 2:**
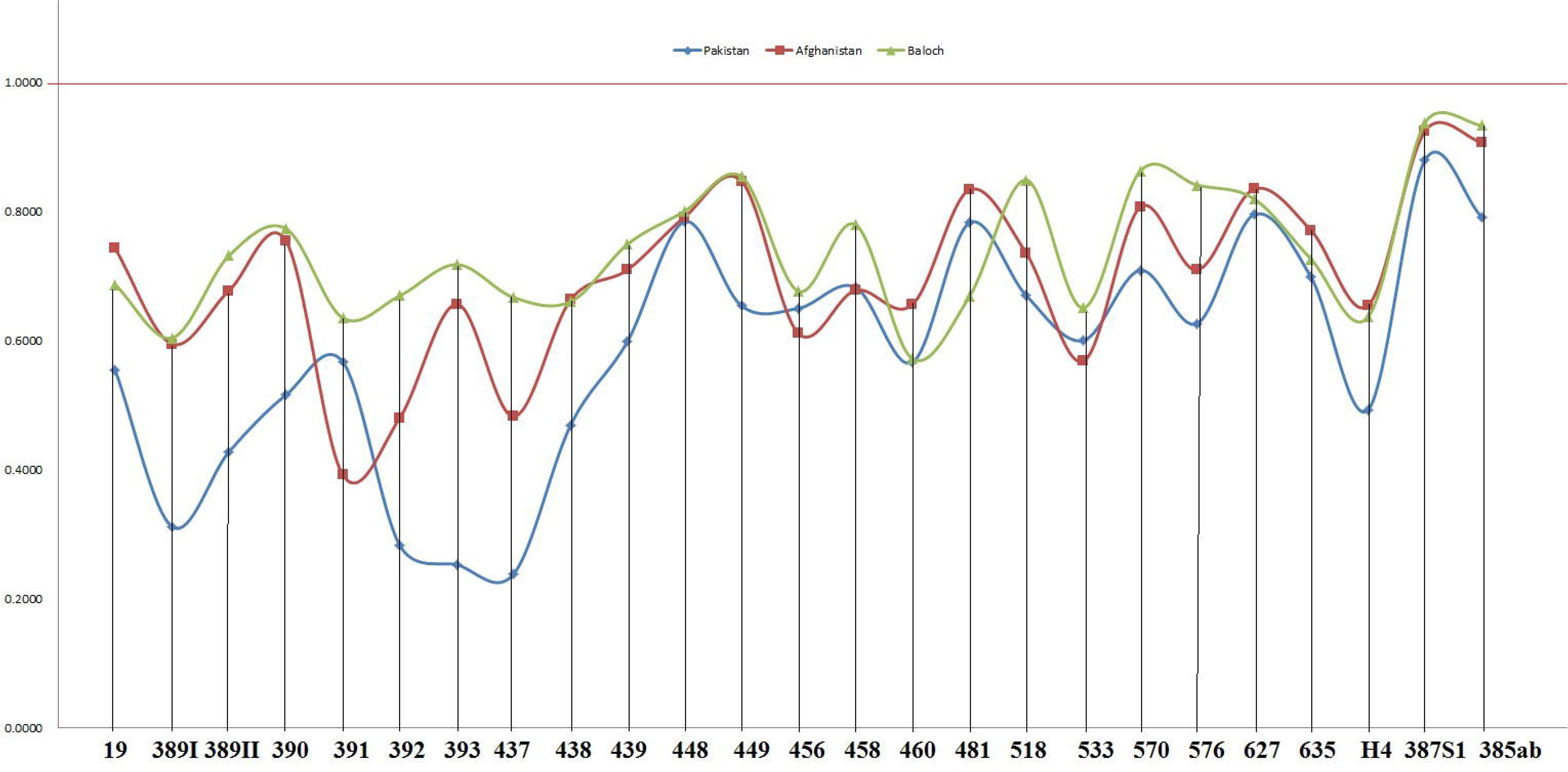
Neighbor-joining tree based on the F*st* values between the Baloch population of Pakistan and Hazara populations across the Durand line with reference populations.

### 2.3 Inference of ancestry based on Y STRs

The Y haplogroups were predicted using the online Y-haplogroup predictor software (http://www.nevgen.org/). C2 (previously known as C3-Star cluster) was the most frequent haplogroup in Pakistani and Afghan Hazaras.

The median-joining network of haplotypes (**Figure 3**) showed a bulky central star-like cluster which represents predicated haplogroup M217 and another big cluster representing haplogroup M420 and comprises many of the identical or highly similar haplotypes. These types of features are usually inferred as past male-lineage expansions^35^. Star-like features of haplotypes comprising haplogroup M217 (C2) have been reported previously in Hazara, Mongol, and Kazakh populations^27,33,36^. An explanation about its origin in Mongolia was about ~ 1,000 years ago ^36^. The frequency of R haplogroup in the Baloch population is 36.03%, 22.22% in Pakistani Hazara, and 21.15% in Afghani Hazara. This haplogroup originated in north Asia about 27,000 years ago (http://isogg.org/tree/index.html). R is one of the most frequent haplogroups in Europe, with its branches reaching 80% of the population in some regions. One branch is believed to have originated in the Kurgan culture, known to be the first speakers of the Indo-European languages and responsible for the domestication of the horse^37^. From somewhere in Central Asia, some descendants of the man carrying the M207 mutation on the Y chromosome headed south to arrive in India about 10,000 years ago^38^. This is one of the frequent haplogroups in Pakistan and North India. In the Baloch population frequency of haplogroup L1 is 22.5% and 1.53% in Afghani Hazara. In sub-continental populations its frequency is about 7-15%^39,40^. Genetic studies suggest that this may be one of the original haplogroups of the creators of Indus Valley Civilization^41,42^. The frequency of L1 is about 28% in Pakistan and Baluchistan, from where the agricultural creators of this civilization emerged^43^. The origins of this haplogroup can be traced to the rugged and mountainous Pamir Knot region in Tajikistan^38^.

**Figure 3:**
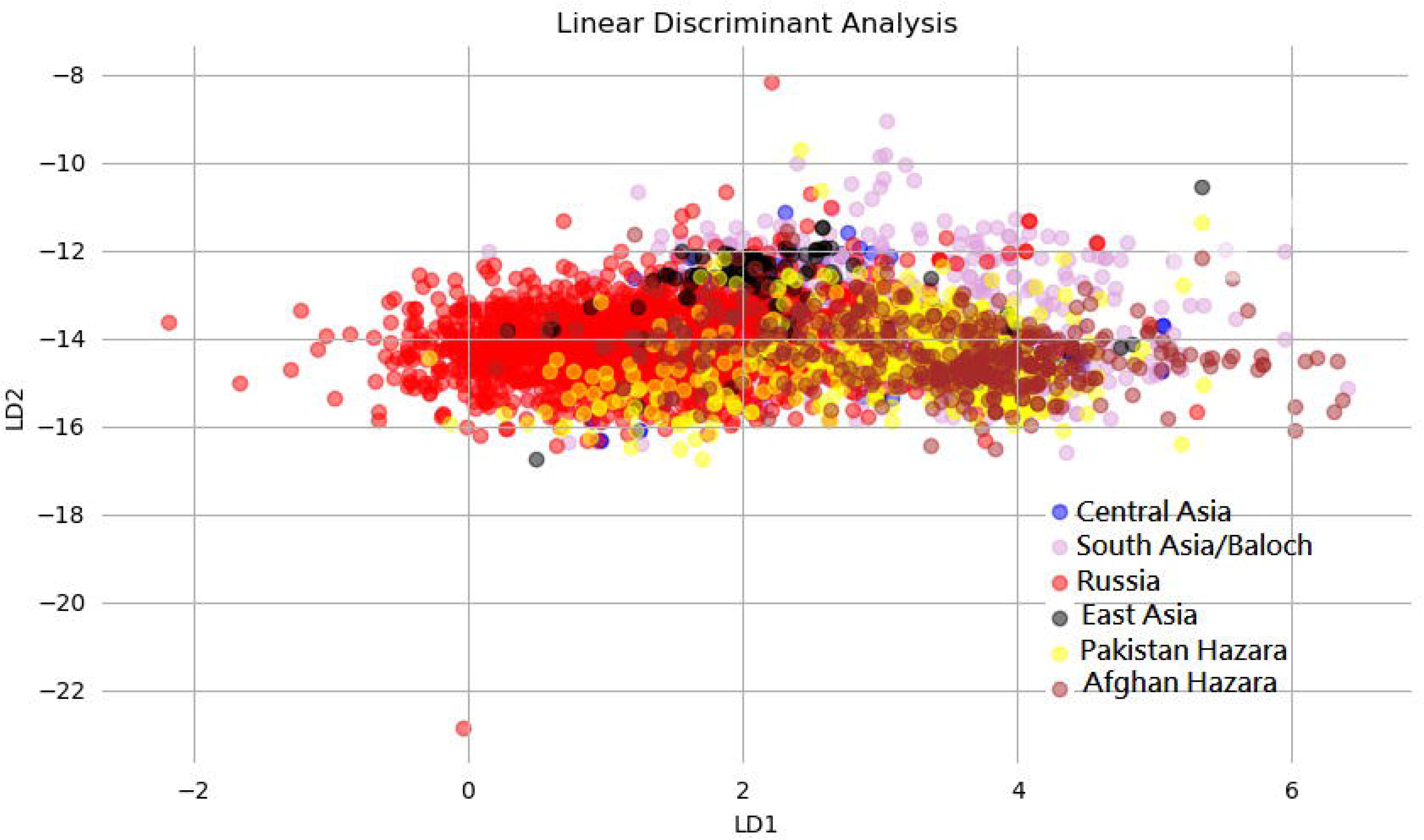
The median-joining network of the Baloch population of Pakistan and Hazara populations across the Durand line based on 20 Y STRs.

In an earlier study^36^, the star-cluster (C3) profile for DYS389I-DYS389b-DYS390-DYS391-DYS392-DYS393-DYS388-DYS425-DYS426-DYS434-DYS435-DYS436-DYS437-DYS438-DYS439 was 10-16-25-10-11-13-14-12-11-11-11-12-8-10-10. In present study mostly occurring haplotype for loci DYS19-DYS389I-DYS389II-DYS390-DYS391-DYS392-DYS393-DYS437-DYS438-DYS439 was 15-13-29-24-10-11-13-14-11-12 which repeated itself in 43 individuals while 14-13-29-24-8-11-13-14-11-11 repeated in 9 individuals and 15-13-29-24-11-11-13-14-11-12 repeated in 8 individuals in Pakistani Hazara population while in Afghani Hazara 16-13-29-25-10-11-13-14-10-10, 15-13-29-24-10-11-13-14-11-12, 14-12-28-23-10-11-12-15-9-11, 14-13-29-24-11-13-12-15-12-12 and 15-14-32-25-11-11-13-14-9-10 haplotypes were repeated in 30, 17, 15, 12 and 11 individuals, respectively. The occurrence of these haplotypes were previously observed in Mongols and Kazakhs^35^. Allelic ranges of Kazak^35^ population from Kazakhstan Central Asia were similar while Mongol population from Inner Mongolia were almost similar on above mentioned 10 Y STRs. In our earlier study^31^, results showed that Hazaras have a close genetic affinity with Turkic-speaking (Kazakh, Kyrgyz and Uyghur) and Mongolian people. Admixture and outgroup findings further clarified that Hazara have 57.8% gene pool from Mongolians.

Here we also speculated a hypothesis that is based on hearsay that Hazaras living in Pakistan are more conserved and they only mate with the Hazaras while across the Durand line the Hazaras mate with other ethnic groups in Afghanistan. Results of gene diversity/heterozygosity and F-statistics tests are also supporting this hypothesis. According to results, all loci showed more diversity in the Hazara population from Afghanistan when compared with the Hazara population from Pakistan (**Figure 4**). F-statistics test within Hazara populations showed variations at four loci only (DYS393-0.05002, DYS449- 0.01694, DYF387S1- 0.00662 and DYS385a/b- 0.00004) (**Supplementary Table 4**). These variations may be the sampling effect, population diversity, or maybe geographical boundaries. LDA is a transformation technique which is commonly used to understand genome diversity and was performed on the Hazara population, Central Asian, South Asian including the Baloch population, East Asian, and Russian population samples to explore their genetic homology. **Figure 5** shows all individual samples plotted on the two LDA factors (F1 and F2). LDA Plot showed the association of the Hazara population with East and Central Asian populations.

**Figure 4:**
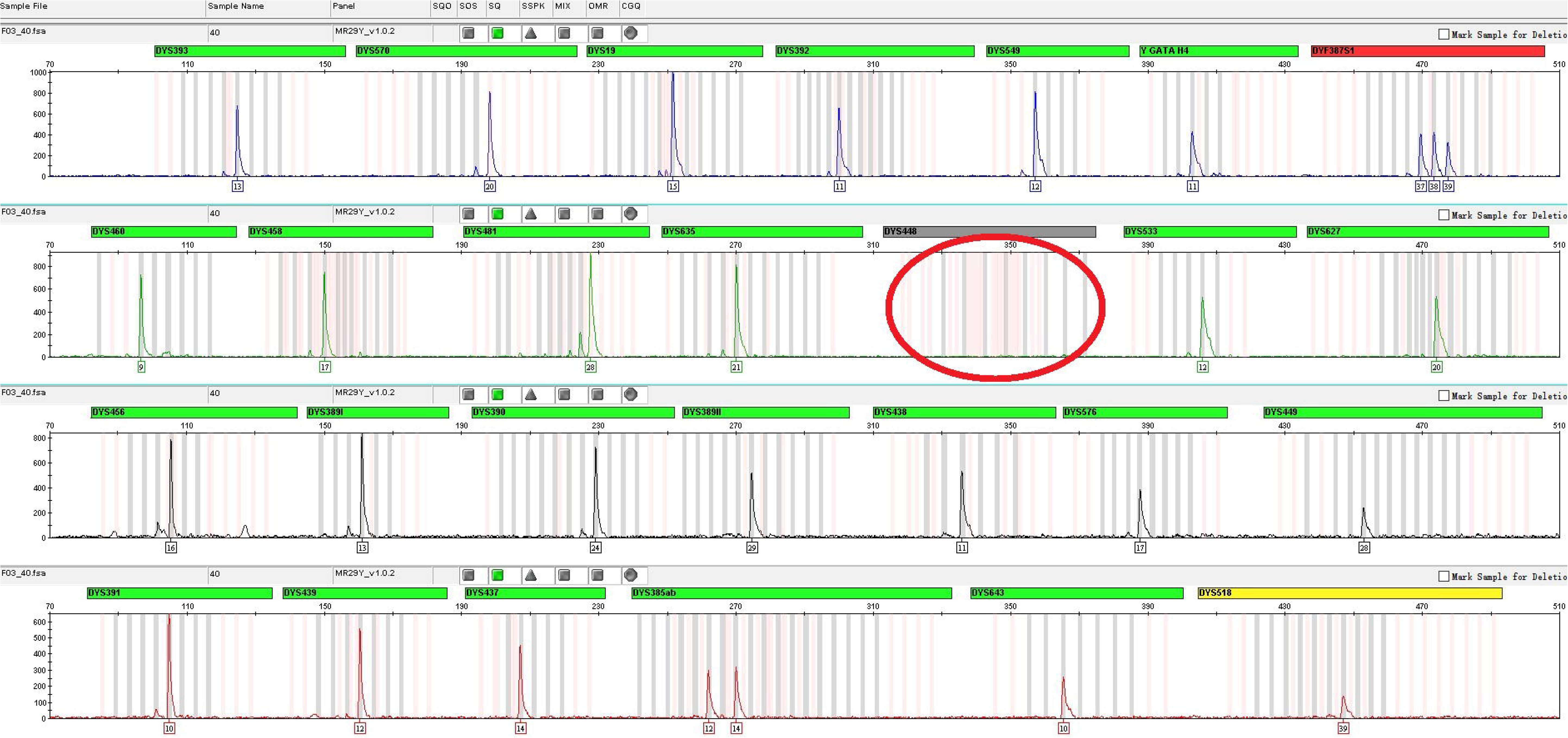
Heterozygosity scattered plot for three populations

**Figure 5:**
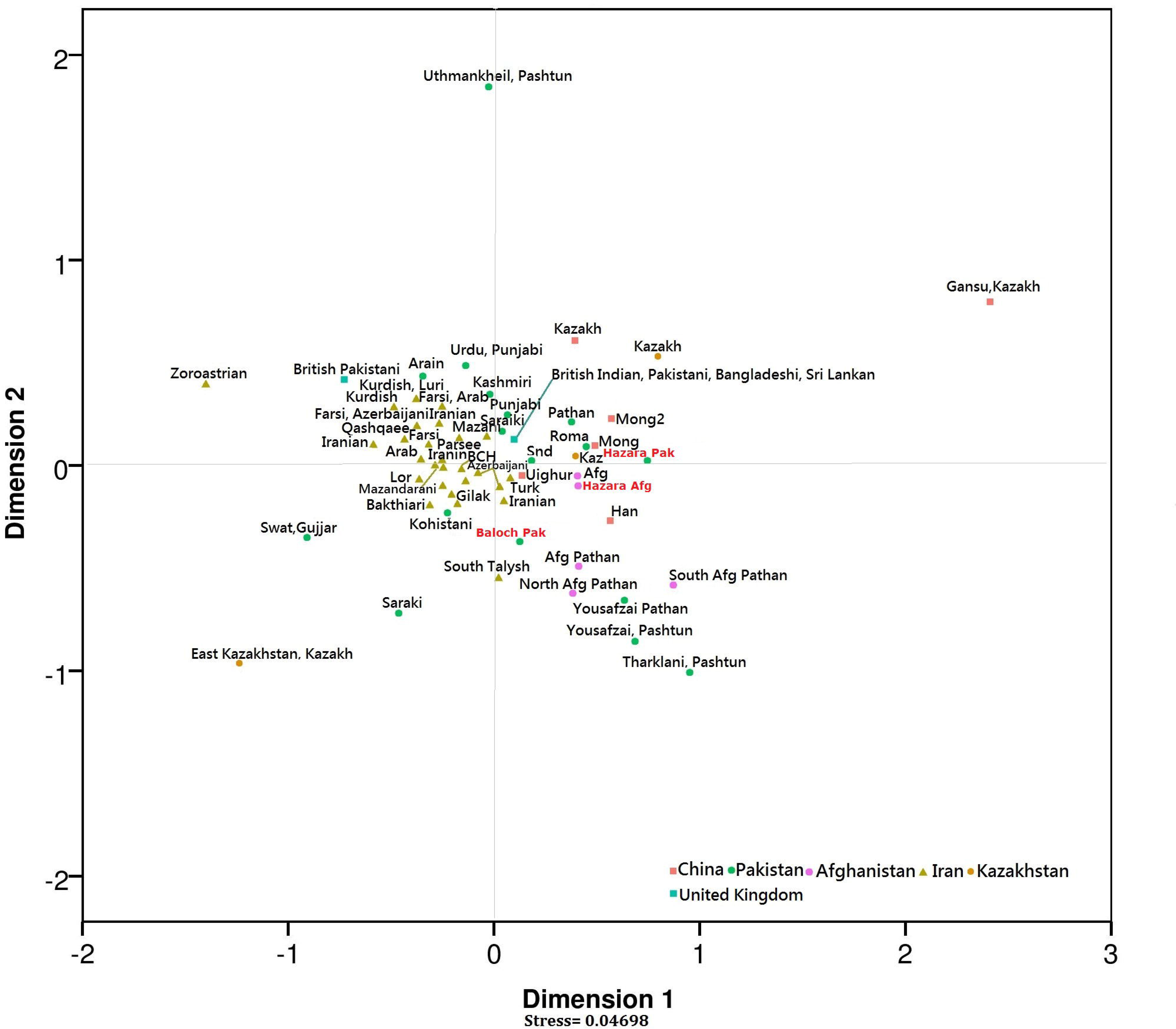
LDA Analysis between the Baloch population of Pakistan and Hazara populations across the Durand line, Central Asia, South Asia, Russia, and East Asian populations.

### 2.4 Physical characterization of DYS448 deletions

By using the Yfiler plus kit, we have observed the null allele at DYS448 in 29 individuals in the Hazara population from Afghanistan (**Figure 6**). Certain factors can cause the phenomena of null alleles and these are deletions within the target region, primer binding sites problem that destabilize hybridization of at least one of the primers flanking the target region ^44–47^. This phenomenon was previously reported, in which other commercial kits were used ^48–53^. The current population study represents the highest frequencies of the null allele at DYS448 when compared with the previously reported population to date (**Table 3**). The core repeat motif of the DYS448 locus is the hexanucleotide repeat AGAGAT^54^. DYS448 has two polymorphic domains separated by an invariant 42-bp region.

**Figure 6:**
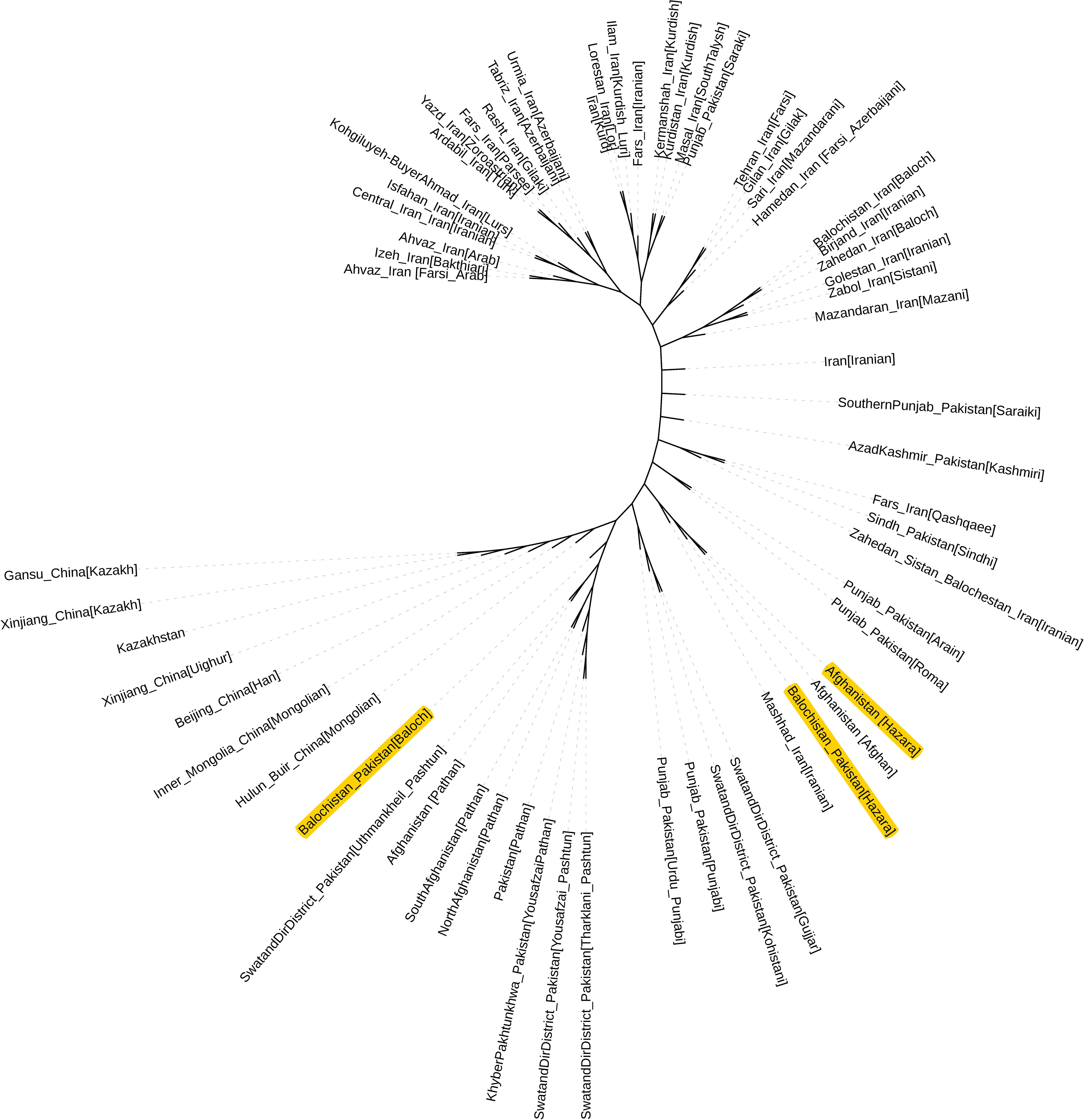
Electropherogram of an individual showing null type at DYS448.

**Table 3:**
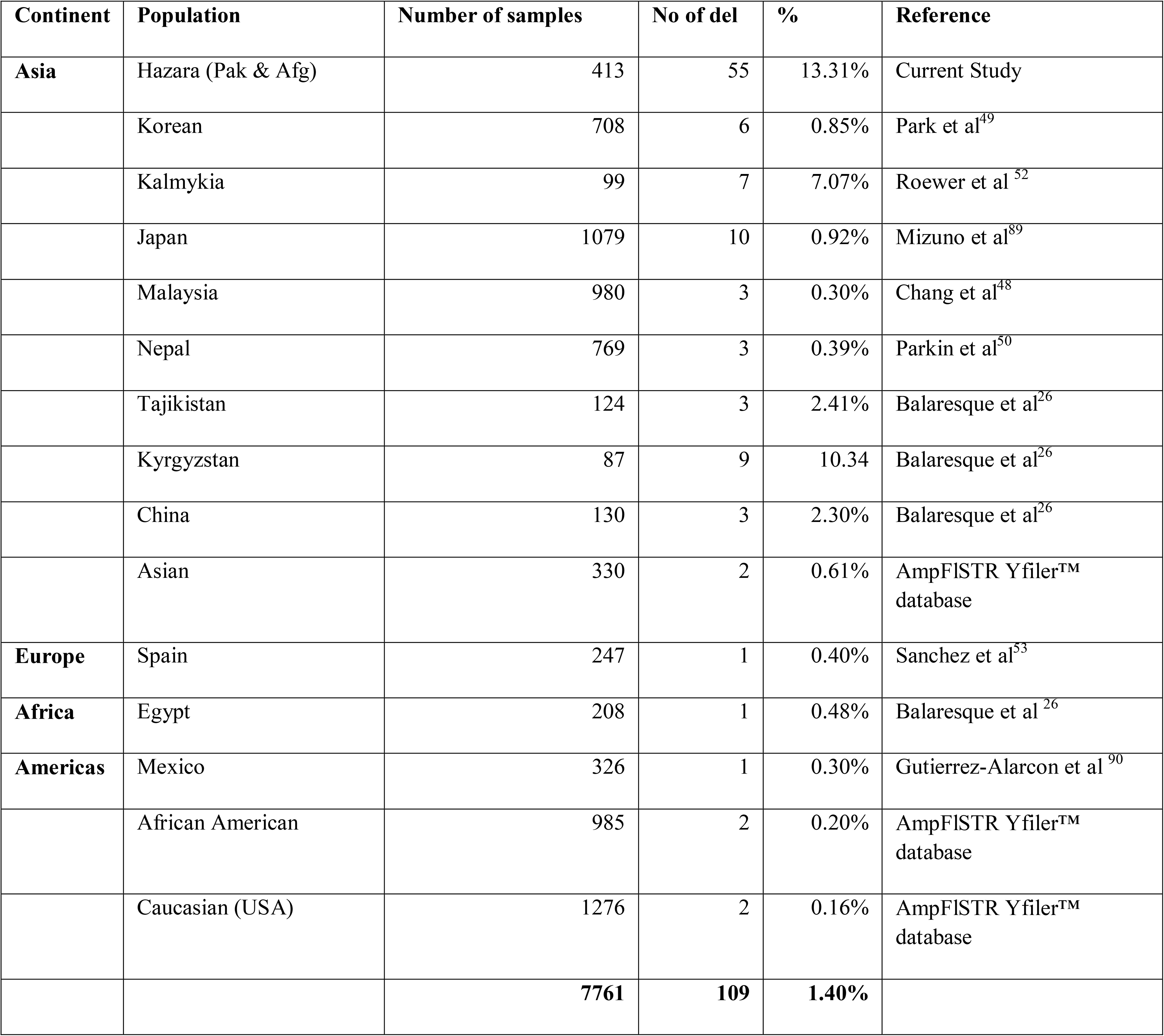
Frequencies of the null allele at DYS448 in various ethnic groups across continents

We have observed 29 null alleles among these, long deletions were covering at a minimum the N42 region and the core AGAGAT repeats downstream, and small deletions encompassing upstream repeats as well (all alignments were based on allele 20). Observed null alleles at locus DYS448 in 29 individuals from the Hazara population of Afghanistan, which were later confirmed with the GoldenEye Y20 System kit were successfully amplified using self-designed primers and sequenced (**Supplementary Table 5**) which were submitted to genbank under accession numbers MN623385 to MN623413. Overall we have observed 55 null alleles at DYS448 in the Hazara population from Pakistan and Afghanistan. Interestingly, all individuals (55) who showed deletion at DYS448 belongs to haplogroup C2 which is most frequent haplogroup in Mongol and Kazakh populations. This high frequency of allele drop-out / mutation is DYS448 in Hazara population from Pakistan and Afghanistan strongly support the evidence that they have Kazakh and Mongol origin. Whole genome or Y Chromosomal sequencing is required to get more insight of this polymorphism. The frequency of the null allele at DYS448 is more frequent in Asia more specifically in East and Central Asia when compared to the rest of the world^26,49^. The commercial companies should pay special attention while designing DYS448 primers.

### 2.5 Concluding Remarks

Finally, our study demonstrates that the Yfiler plus kit detects high haplotype diversity in Baloch population from Pakistan and Hazara populations from across the Durand line (Pakistan and Afghanistan) of which two (Baloch and Afghani Hazara) were not previously studied at Yfiler plus STR loci, which in general makes it suitable for forensic casework in these groups. The recent inclusion of these data in the YHRD allows widespread use for forensic and other purposes.

## 3. MATERIALS AND METHODS

### 3.1 Samples

A total of 524 blood samples were collected, in which 111 Balochi individuals from Baluchistan Pakistan, 153 from Hazara Town Quetta, Baluchistan Pakistan (Participants were part of an earlier study ^27^ and were agreed to the secondary use of their DNA samples), and 260 from Bamyan, Afghanistan. All participants who were included in this study were unrelated individuals of at least three generations. All participants gave their informed consent either orally and with thumbprint (in case they could not write) or in writing after the study aims and procedures were carefully explained to them. This collaborative study was approved by the ethical review boards of China Medical University, Shenyang, Liaoning Province, People’s Republic of China (2019/067-P), University of Health Sciences Lahore Pakistan (2017-CMU-1/14), and Ministry of Public Health, Forensic Medicine Directorate, Kabul, Afghanistan (FC-2017-02). All the experimental procedures were performed in accordance with the standards of the Declaration of Helsinki.

### 3.2. DNA extraction

Axygen AxyPrep Blood Genomic DNA Miniprep Kit was used to extract genomic DNA according to the manufacturer’s protocol (Axygen Biosciences; CA, USA).

### 3.3 PCR Amplification

DNA was amplified using Yfiler Plus PCR Amplification Kit (Thermo Fisher Scientific) PCR amplification was carried out using the Applied Biosystems GeneAmp PCR System 9700 thermal cyclers. PCR amplifications were performed as recommended by the manufacturer, although using half of the recommended reaction volume (12.5 μl).

### 3.4. 27Y-STRs genotyping

After successful PCR amplification, The PCR products were analyzed by using an 8 capillary ABI 3500 DNA Genetic Analyzer with POP-4 polymer (Life Technologies) according to the manufacturer’s protocol. GeneMapper Software version 4.0 (Life Technologies) was used for the genotype assignment. DNA typing was performed according to the manufacturer’s protocol by using the locus panel and allele bins supplied by the manufacturer and allele designations corresponding with the allelic ladder supplied by the manufacturer. Genotype nomenclature was based on the recommendations of the International Society for Forensic Genetics ^55^.

### 3.5. Confirmation of Null DYS 448

For the confirmation of samples that showed no allele call at DYS448, they were re-amplified by using the Goldeneye 20Y amplification kit (Goldeneye Technology Ltd.). After confirmed with two different kits (Yfiler Plus and GoldenEye 20Y), these samples were amplified and sequenced as described elsewhere ^27^.

### 3.6. Quality control

Our laboratory has participated and passed the YHRD quality assurance exercise 2015. Haplotype data were already made accessible via the Y-chromosome Haplotype Reference Database (YHRD) under accession number YA004595 (Balochi) in 61st release on dated 2019 June 24, YA004312-2 (Hazara Pakistan) and YA004503 (Hazara Afghanistan) in 59th release on dated 2018 November 01. 29 sequenced samples at null allele call at DYS448 were also submitted to genbank under accession numbers MN623385 to MN623413 on dated 2019 october 28.

### 3.7. Statistical analysis

Allelic and haplotype frequencies were computed by direct counting method and haplotype diversity (HD) was calculated according to:

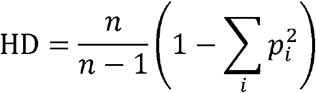

where *n* is the male population size and *p_i_* is the frequency of *i*th haplotype. Discrimination capacity (DC) was calculated as the ratio of unique haplotypes in the samples. Match probabilities (MP) were calculated as Σ P*i*^2^, where P*i* is the frequency of the *i*-th haplotype. Genetic distances were evaluated using the Rst^56^ and Fst^57–59^ statistic, between reference populations and currently studied populations on overlapping STRs (DYS19, DYS389I, DYS389II, DYS390, DYS391, DYS392, DYS393, DYS437, DYS438, DYS439, DYS448, DYS456, DYS458, DYS635, and Y_GATA_H4) were calculated by using Arlequin Software v3.5^60^. We calculated both Rst and Fst values because in the generalized stepwise mutation model, Rst offers relatively unbiased evaluations of migration rates and times of population divergence while on other hand Fst tends to show too much population similarity, predominantly when migration rates are low or divergence times are long^56^. Reduced dimensionality spatial representation of the populations was performed based on Rst values using multi-dimensional scaling (MDS) with IBM SPSS Statistics for Windows, Version 23.0 (IBM Corp., Armonk, NY, USA). Heatmaps were generated using Rst and Fst values were generated using R program V3.4.1 platform with the help of a ggplot2 module.

#### Phylogenetic analysis

A neighbor-joining phylogenetic tree was constructed for the Hazara and the reference populations based on a distance matrix of Fst using the Mega7 software^61^. We also predicted Y-SNP haplogroups in the samples from Y STR haplotypes (Yfiler STRs) using the Y-DNA Haplogroup Predictor NEVGEN (http://www.nevgen.org). We have used FTDNA order for 17 Y STRs (Yfiler loci). The microvariant alleles were truncated to the next lowest integer value since values in the database were treated similarly. Any haplotypes which have null alleles or duplication variants in the Baloch or Hazara population from Pakistan or Afghanistan were excluded from the analysis. The results of NEVGEN were cross checked with Athey’s Haplogroup Predictor (http://www.hprg.com/hapest5/index.html).

#### Linear discriminant analysis

R program V3.4.1 platform with the help of a ggplot2 module was used to perform linear discriminant analysis (LDA) for Hazara (Pakistan), Hazara (Afghanistan), Central Asia, East Asia, the Middle East, and Southwest Asian (Baloch) samples ^62^ on overlapping (DYS19, DYS389I, DYS389II, DYS390, DYS391, DYS392, DYS393, DYS437, DYS438, DYS439, DYS448, DYS456, DYS458, DYS635, and Y_GATA_H4) STRs. The multi-copy marker like (DYS385ab) and haplotypes that have null alleles or duplication variants in the Baloch or Hazara population or any of the reference populations were excluded from the analysis. For DYS389I and DYS389II, we have subtracted DYS389I from DYS389II and used DYS389II-I for analysis.

#### The median-joining network

To define the genetic relationships among Balochi and Hazara individuals for 20 Y STRs (DYS19, DYS389II-I, DYS390, DYS391, DYS392, DYS393, DYS437, DYS438, DYS448, DYS456, DYS458, DYS635, Y_GATA_H4, DYS549, DYS460, DYS481, DYS533, DYS570, DYS576, DYS627), we used the stepwise mutation model and Median Joining-Maximum Parsimony algorithm ^63^ by using the program Network 5 as described at the Fluxus Engineering website (http://www.fluxus-engineering.com), and the weighting criteria for Y-STRs following ^27^ Any haplotypes which have null alleles or duplication variants in the Baloch or Hazara population from Pakistan or Afghanistan were excluded from the analysis.

## Supporting information

Supplementary Table 4: F-statistics analysis between Hazara population from Pakistan and Afghanistan

Supplementary Table 5: Sequence in the relevant flanking and repeat region of the DYS448 locus for null alleles.

Supplementary Figure 1: Heatmap generated using Rst and Fst values.

Supplementary Figure 1: Heatmap generated using Rst and Fst values.

Supplementary Table 1: Raw genotypic data of 3 ethnic groups typed with Yfiler plus

Supplementary Table 2: Allele Frequencies and Forensic Parameters 3 ethnic groups

Supplementary Table 3: Pairwise Rst and Fst values between 3 ethnic groups and other reference populations

## AUTHOR CONTRIBUTION

J.L. and A.A. designed this study. A.A., A.R., and S.N. and M.R., collected the samples. A.A. experimented and wrote the manuscript. A.A., J.L., A.R., S.N., R.A., S.W., and C.W., analyzed the results. A.A., and J.L., modified the manuscript. All authors reviewed the manuscript.

## COMPLIANCE WITH ETHICAL STANDARDS

The study was approved (2019/067-P) by the ethical review board of China Medical University, Shenyang, Liaoning Province, People’s Republic of China, and in accordance with the standards of the Declaration of Helsinki. All participants who were included in this study were unrelated individuals of at least three generations. All participants gave their informed consent either orally and with thumbprint (in case they could not write) or in writing after the study aims and procedures were carefully explained to them.

## ACKNOWLEDGMENTS

We thank all volunteers who provided material and data for this project, especially Guanglin He and Abul-Hasan Fawad. This study was financially supported by the China Medical University postdoctoral research grant (100/1210619014).

## Electronic Supplementary Materials (ESM)

**Supplementary Figure 1:** Heatmap generated using Rst and Fst values.

**Supplementary Table 1:** Raw genotypic data of 3 ethnic groups typed with Yfiler plus

**Supplementary Table 2:** Allele Frequencies and Forensic Parameters 3 ethnic groups

**Supplementary Table 3:** Pairwise R*st* and F*st* values between 3 ethnic groups and other reference populations

**Supplementary Table 4**: F-statistics analysis between Hazara population from Pakistan and Afghanistan

**Supplementary Table 5:** Sequence in the relevant flanking and repeat region of the DYS448 locus for null alleles.

