## Supplementary figures and images for "Forensic features and genetic legacy of the Baloch population of Pakistan and the Hazara population across Durand-line revealed by Y chromosomal STRs"

### Supplementary Figure 1: Heatmap generated using Rst and Fst values.

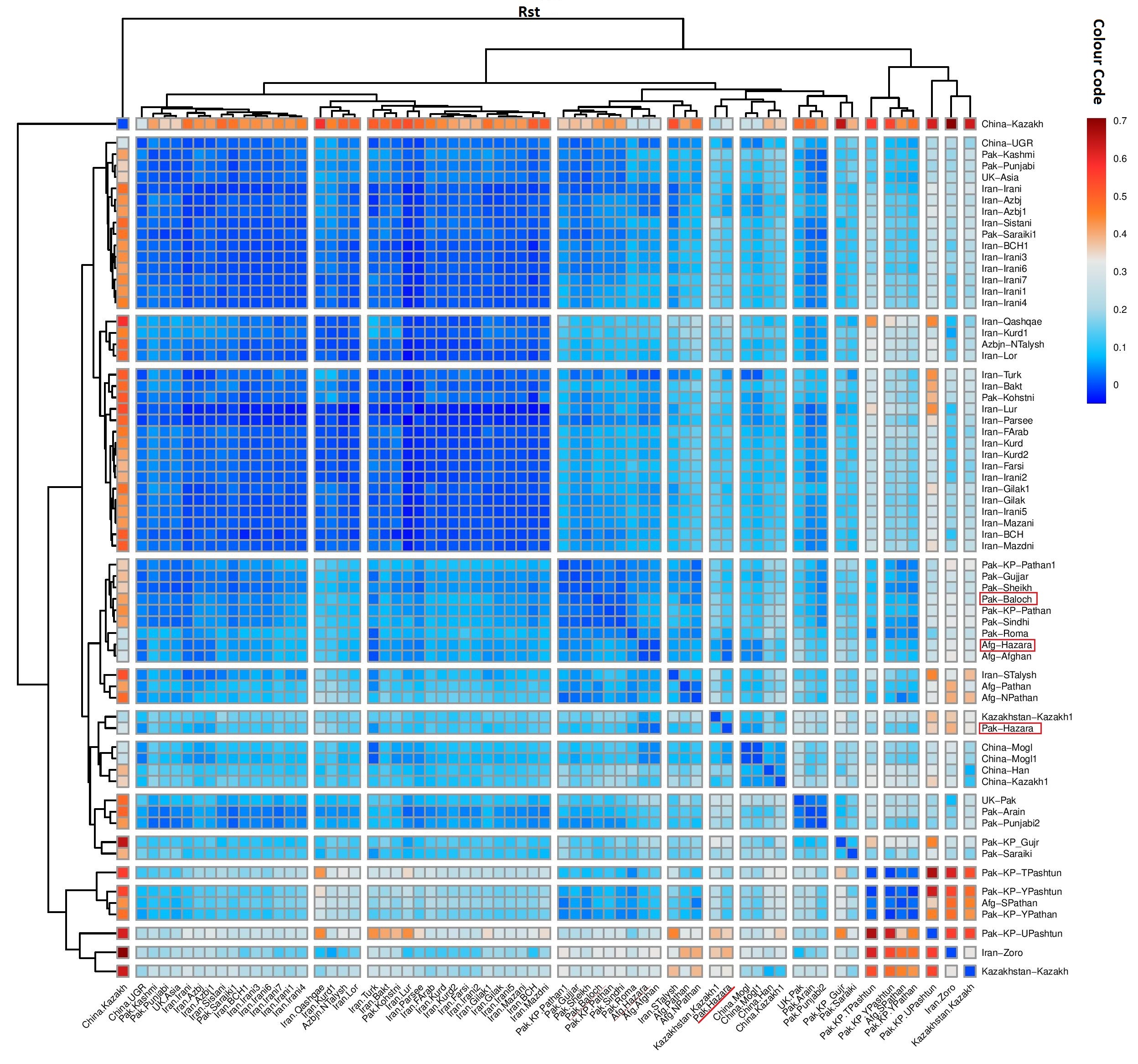

### Supplementary Figure 1: Heatmap generated using Rst and Fst values.

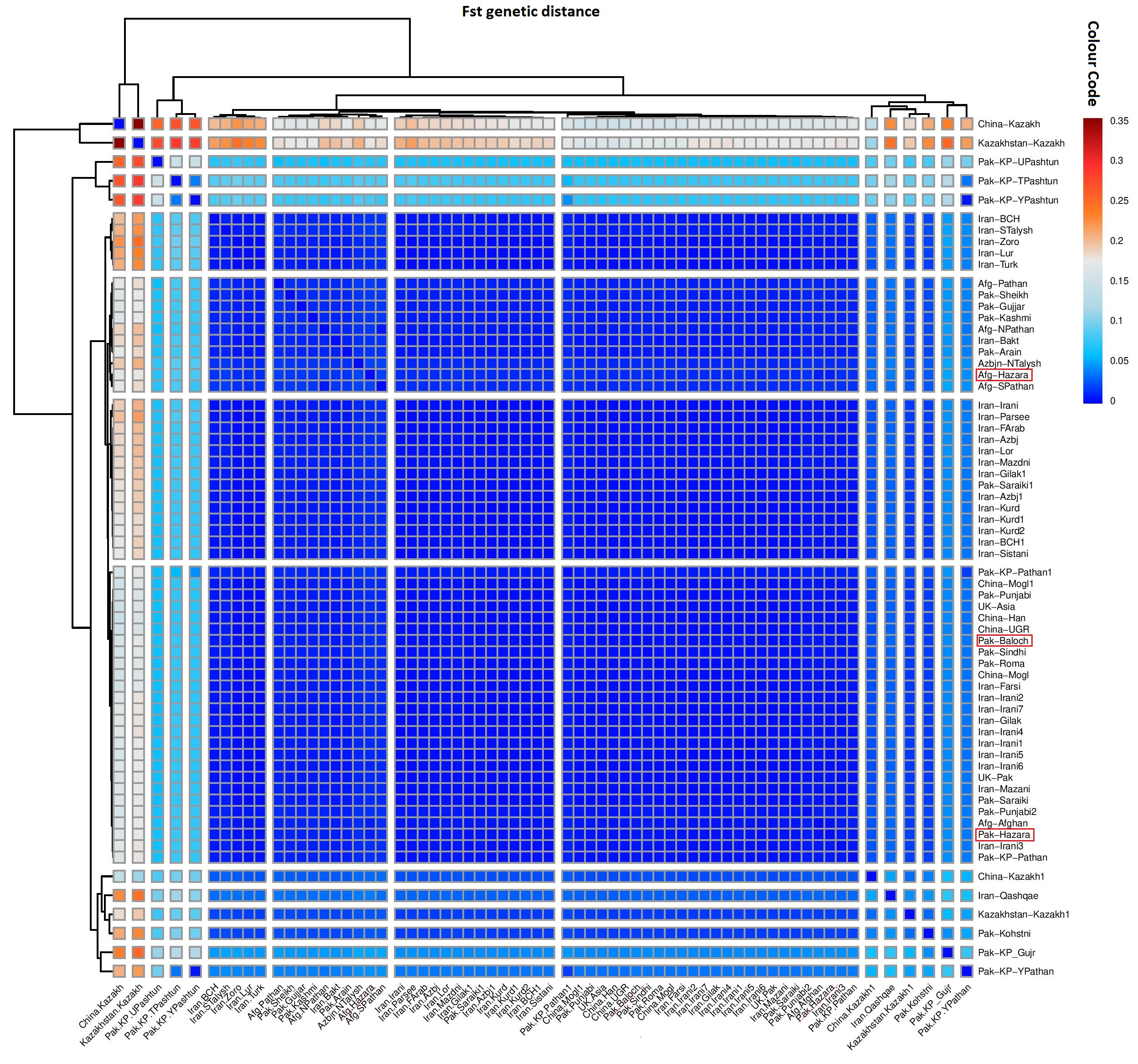
